# Gene expression patterns associated with neurological disease in HIV infection

**DOI:** 10.1101/096172

**Authors:** Pietro Paolo Sanna, Vez Repunte-Canonigo, Eliezer Masliah, Celine Lefebvre

**Author notes:** Corresponding Author:* Pietro Paolo Sanna, MD, The Scripps Research Institute, 10550 North Torrey Pines Road, La Jolla, CA 92037-1000. Abbreviations: Asymptomatic neurocognitive impairment (ANI), Combination antiretroviral therapy (cART), Gene Set Enrichment Analysis (GSEA), HIV-associated neurological disease (HAND), HIV encephalitis (HIVE), Interferon (IFN), NeuroAIDS Tissue Consortium (NNTC), Neurocognitive impairments (NCI).

## Abstract

To provide new insight into the pathogenesis of neurocognitive impairments (NCI) in HIV infection, we used the Gene Set Enrichment Analysis (GSEA) algorithm to analyze pathway dysregulations in gene expression profiles of HIV-infected patients with or without NCI and HIV encephalitis (HIVE). While HIVE was characterized by widespread inflammation and tissue damage, gene expression evidence of induction of interferon (IFN), cytokines and tissue injury was apparent in all brain regions studied before the emergence of NCI. Various degrees of white matter changes were present in all HIV-infected subjects and were the primary manifestation in patients with NCI in the absence of HIVE. The latter showed a distinct pattern of immune activation with induction of chemokines, cytokines, β-defensins, and limited IFN induction.

Altogether results indicate that significant neuroinflammation and neuronal suffering precede NCI. Patients with NCI without HIVE showed a predominantly white matter dysfunction with a distinct pattern of immune activation.

## Introduction

While the prevalence of severe HIV-associated dementia (HAD) has decreased since the introduction of combination antiretroviral therapy (cART), milder and chronic forms of neurocognitive impairment (NCI) including asymptomatic neurocognitive impairment (ANI) and HIV-associated neurocognitive disorders (HAND) as well as HIV-associated major depressive disorder remain high (1–7). HIV encephalitis (HIVE) is considered to be the main neuropathological substrate of HAD (8–10). NCI in the setting of cART is associated with synaptodendritic degeneration (7, 11, 12). While the brain represents a sanctuary where HIV can persist due to suboptimal penetration of antiretroviral drugs (13), various studies highlighted the occurrence of NCI even in the setting of viral suppression (14, 15). Chronic neuroinflammation is believed to drive neurodegeneration in cART-era HAND (7, 9, 16, 17). However, the pathogenic mechanisms behind HAND remain unclear.

To identify gene expression correlates of neurological disease progression in HIV, we analyzed pathway dysregulations in brain regions of patients in the National NeuroAIDS Tissue Consortium (NNTC) gene expression profile dataset. The NNTC dataset consists of samples from 3 different brain regions (white matter, basal ganglia, prefrontal cortex) of control and HIV-infected patients with or without NCI and HIVE (18). For pathway analysis we used the Gene Set Enrichment Analysis (GSEA), a computational method to assess whether a priori defined sets of genes show statistically significant differences between biological states (19). GSEA was used in conjunction with gene sets from the Molecular Signatures Database (MSigDb), including canonical pathways in the C2 collection (20).

## Materials and Methods

### Dataset and analysis

Clinical and demographic features of the subjects in the NNTC gene expression dataset used for the study are shown in Supplementary Table 1. Raw data were downloaded from GEO (GSE35864). We filtered out 9 samples based on quality controls (actin3/actin5 ratio, gapdh3/gapdh5 ratio, NUSE (Normalized Unscaled Standard Errors) and RLE (Relative Log Expression) computed with the packages simpleaffy and affyPLM in R). Normalization was done using gcrma (21). We further checked the expression of markers of neurons (RBFOX3) and olygodendrocytes (MBP) to validate the brain region profiled for white matter and prefrontal cortex samples and further excluded 2 samples that had conflicting expression according to their classification (D1 WM and D1 FC).

**Table 1.**
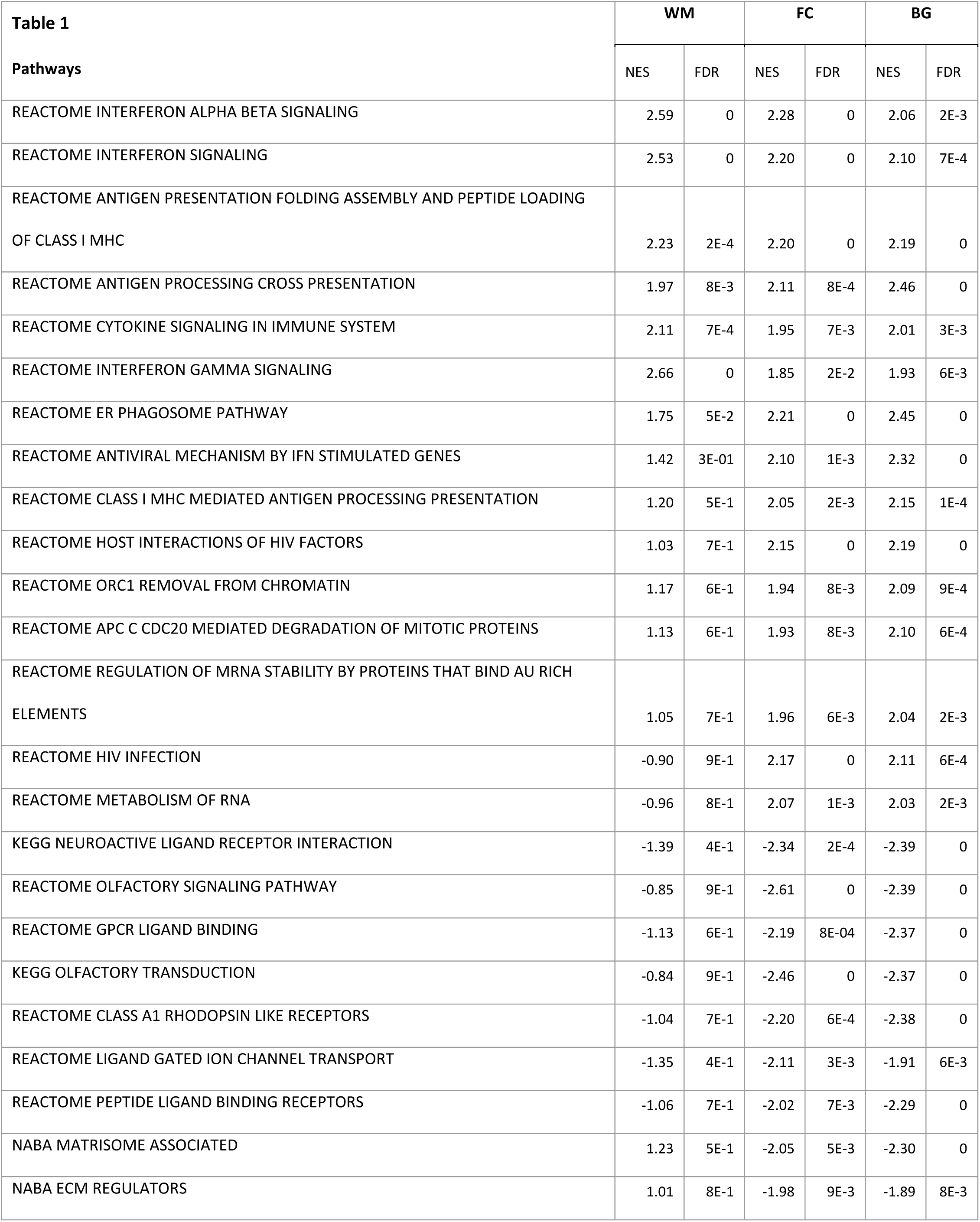
Pathways differentially regulated in multiple brain regions in patients infected with HIV without NCI as compared to uninfected controls.

| Table 1<br>Pathways | WM |  | FC |  | BG |  |
| --- | --- | --- | --- | --- | --- | --- |
|  | NES | FDR | NES | FDR | NES | FDR |
| REACTOME INTERFERON ALPHA BETA SIGNALING | 2.59 | 0 | 2.28 | 0 | 2.06 | 2E-3 |
| REACTOME INTERFERON SIGNALING | 2.53 | 0 | 2.20 | 0 | 2.10 | 7E-4 |
| REACTOME ANTIGEN PRESENTATION FOLDING ASSEMBLY AND PEPTIDE LOADING OF CLASS I MHC | 2.23 | 2E-4 | 2.20 | 0 | 2.19 | 0 |
| REACTOME ANTIGEN PROCESSING CROSS PRESENTATION | 1.97 | 8E-3 | 2.11 | 8E-4 | 2.46 | 0 |
| REACTOME CYTOKINE SIGNALING IN IMMUNE SYSTEM | 2.11 | 7E-4 | 1.95 | 7E-3 | 2.01 | 3E-3 |
| REACTOME INTERFERON GAMMA SIGNALING | 2.66 | 0 | 1.85 | 2E-2 | 1.93 | 6E-3 |
| REACTOME ER PHAGOSOME PATHWAY | 1.75 | 5E-2 | 2.21 | 0 | 2.45 | 0 |
| REACTOME ANTIVIRAL MECHANISM BY IFN STIMULATED GENES | 1.42 | 3E-01 | 2.10 | 1E-3 | 2.32 | 0 |
| REACTOME CLASS I MHC MEDIATED ANTIGEN PROCESSING PRESENTATION | 1.20 | 5E-1 | 2.05 | 2E-3 | 2.15 | 1E-4 |
| REACTOME HOST INTERACTIONS OF HIV FACTORS | 1.03 | 7E-1 | 2.15 | 0 | 2.19 | 0 |
| REACTOME ORC1 REMOVAL FROM CHROMATIN | 1.17 | 6E-1 | 1.94 | 8E-3 | 2.09 | 9E-4 |
| REACTOME APC C CDC20 MEDIATED DEGRADATION OF MITOTIC PROTEINS | 1.13 | 6E-1 | 1.93 | 8E-3 | 2.10 | 6E-4 |
| REACTOME REGULATION OF MRNA STABILITY BY PROTEINS THAT BIND AU RICH ELEMENTS | 1.05 | 7E-1 | 1.96 | 6E-3 | 2.04 | 2E-3 |
| REACTOME HIV INFECTION | -0.90 | 9E-1 | 2.17 | 0 | 2.11 | 6E-4 |
| REACTOME METABOLISM OF RNA | -0.96 | 8E-1 | 2.07 | 1E-3 | 2.03 | 2E-3 |
| KEGG NEUROACTIVE LIGAND RECEPTOR INTERACTION | -1.39 | 4E-1 | -2.34 | 2E-4 | -2.39 | 0 |
| REACTOME OLFACTORY SIGNALING PATHWAY | -0.85 | 9E-1 | -2.61 | 0 | -2.39 | 0 |
| REACTOME GPCR LIGAND BINDING | -1.13 | 6E-1 | -2.19 | 8E-04 | -2.37 | 0 |
| KEGG OLFACTORY TRANSDUCTION | -0.84 | 9E-1 | -2.46 | 0 | -2.37 | 0 |
| REACTOME CLASS A1 RHODOPSIN LIKE RECEPTORS | -1.04 | 7E-1 | -2.20 | 6E-4 | -2.38 | 0 |
| REACTOME LIGAND GATED ION CHANNEL TRANSPORT | -1.35 | 4E-1 | -2.11 | 3E-3 | -1.91 | 6E-3 |
| REACTOME PEPTIDE LIGAND BINDING RECEPTORS | -1.06 | 7E-1 | -2.02 | 7E-3 | -2.29 | 0 |
| NABA MATRISOME ASSOCIATED | 1.23 | 5E-1 | -2.05 | 5E-3 | -2.30 | 0 |
| NABA ECM REGULATORS | 1.01 | 8E-1 | -1.98 | 9E-3 | -1.89 | 8E-3 |

### Pathway analysis

For pathway analysis, we selected one representative probe per gene based on the highest observed coefficient of variation of the probes across the samples. The dataset was interrogated for pathway enrichment using the GSEA algorithm and the canonical pathways from the MSidDb C2 collection (1,237 pathways with at least 10 genes). GSEA was run using 1,000 shuffling of the reference list. Significance was assessed using the False Discovery Rate (FDR) computed as defined in the original GSEA publication for controlling the number of false positives in each GSEA analysis (19). Differential expression was computed using a Welch t-test from the package Class Comparison in R 3.3.1. We defined the pathways commonly differentially regulated in each comparison as the pathways satisfying an FDR < 0.01 in at least 2 of the 3 brain regions for that comparison while pathways specific to one region were defined as pathways satisfying FDR < 0.01 in that region and FDR > 0.25 in the other 2 regions.

### Pathways activity

The activity of a pathway in a sample was computed the following way: we first z-transformed the gene expression profiles to normalize the expression of each gene across samples. We then computed the enrichment score (ES) of a gene set using this z-transformed matrix of expression, as described in the original description of GSEA (19). The ES corresponds to the relative activity of a gene set in a sample as compared to all others. Hence, the samples with the highest ES are the samples with the highest relative expression of the genes belonging to this set among the samples belonging to the gene expression matrix.

### GSEA

Gene set enrichment analysis was implemented in R and follows the method described in (19). Null distribution was obtained by 1,000 shuffling of the reference list. Gene signatures were obtained by ranking the genes according to the sign of the statistics (S) and the p-value (p) of the test with the following metric: - 1 ×sign(S)×log(P,10).

## Results

### Pathway analysis of the NNTC dataset

The NNTC dataset was interrogated for pathway enrichment using the canonical pathways from the MSigDb C2 collection and the GSEA algorithm (19) (See methods). We compared the following four groups: A: controls; B: HIV-infected no NCI no HIVE; C: HIV-infected with NCI no HIVE; D: HIV-infected with NCI and HIVE (Fig. 1). All results are presented in supplementary tables in Supplementary Tables 2–6.

**Figure 1:**
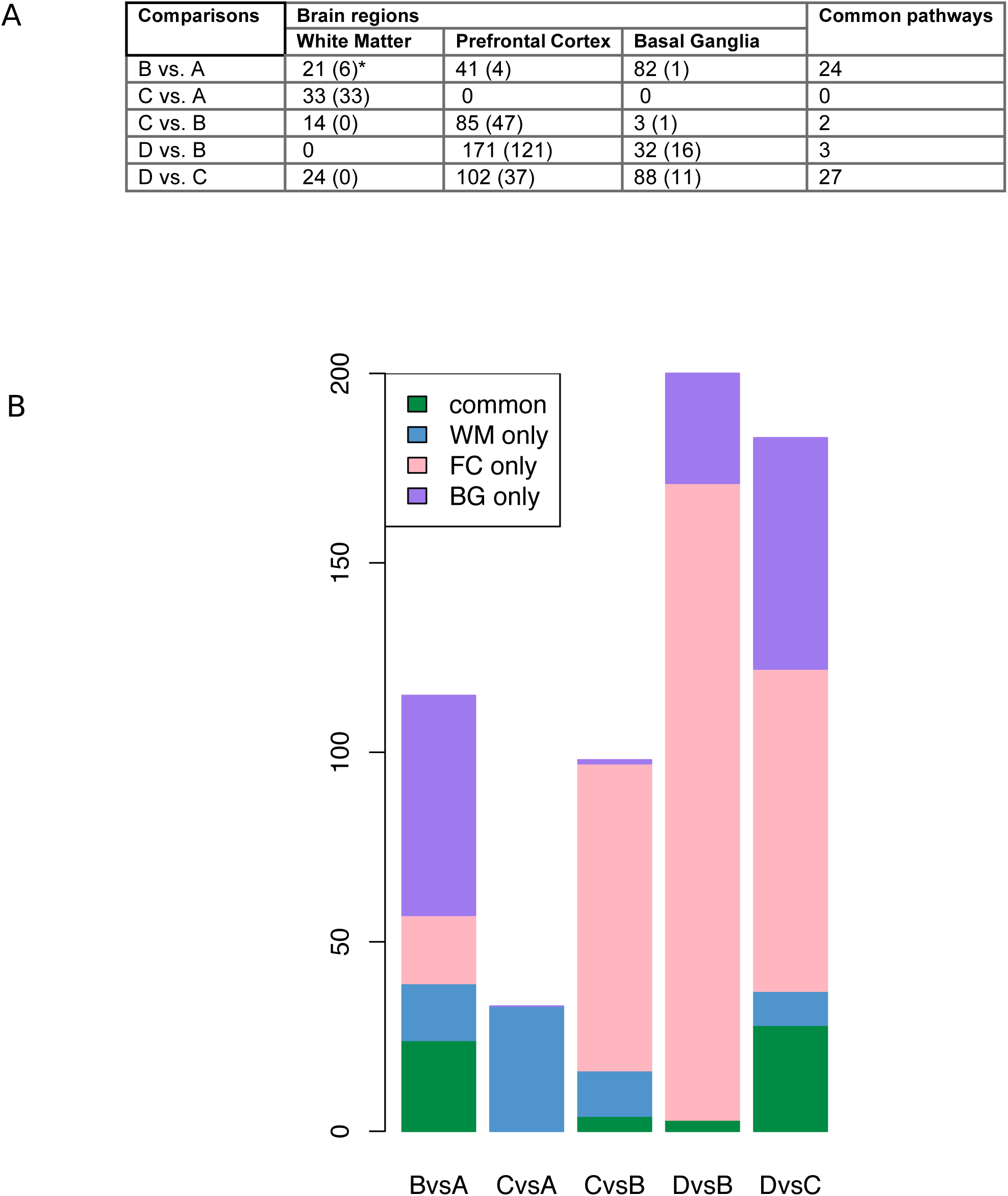
**A) Number of Pathways differentially regulated in each transition in each brain region.** *Numbers in brackets indicate the number of pathways selectively differentially regulated in that region for a particular transition as compared to the other 2 brain regions. **B) Bar plot showing the number of pathways significantly differentially regulated per comparison (FDR < 0.01).** WM=White Matter; FC = Frontal Cortex; BG = Basal Ganglia. Common pathways define pathways significant in at least 2 regions.

### Identification of pathways dysregulated in HIV-infected patients without NCI vs. uninfected controls (B-A comparison)

We identified 24 pathways concordantly differentially regulated in at least 2 brain regions in this transition (Table 1). Genes driving the enrichment (on the left of the leading edge corresponding to the peak of the running enrichment in GSEA as shown in Fig. 2A) were retrieved for each region (Fig. 3). These pathways and genes indicate a significant activation of IFN and cytokine signaling prior to the onset of NCI. Both genes regulated by type I and type II IFN were activated in HIV infection without NCI (Fig. 2, 3). IFN-regulated genes, such as MCH class I genes, were induced in all brain regions of HIV-infected patients without NCI as compared to uninfected controls (Fig. 2, 3).

**Figure 2:**
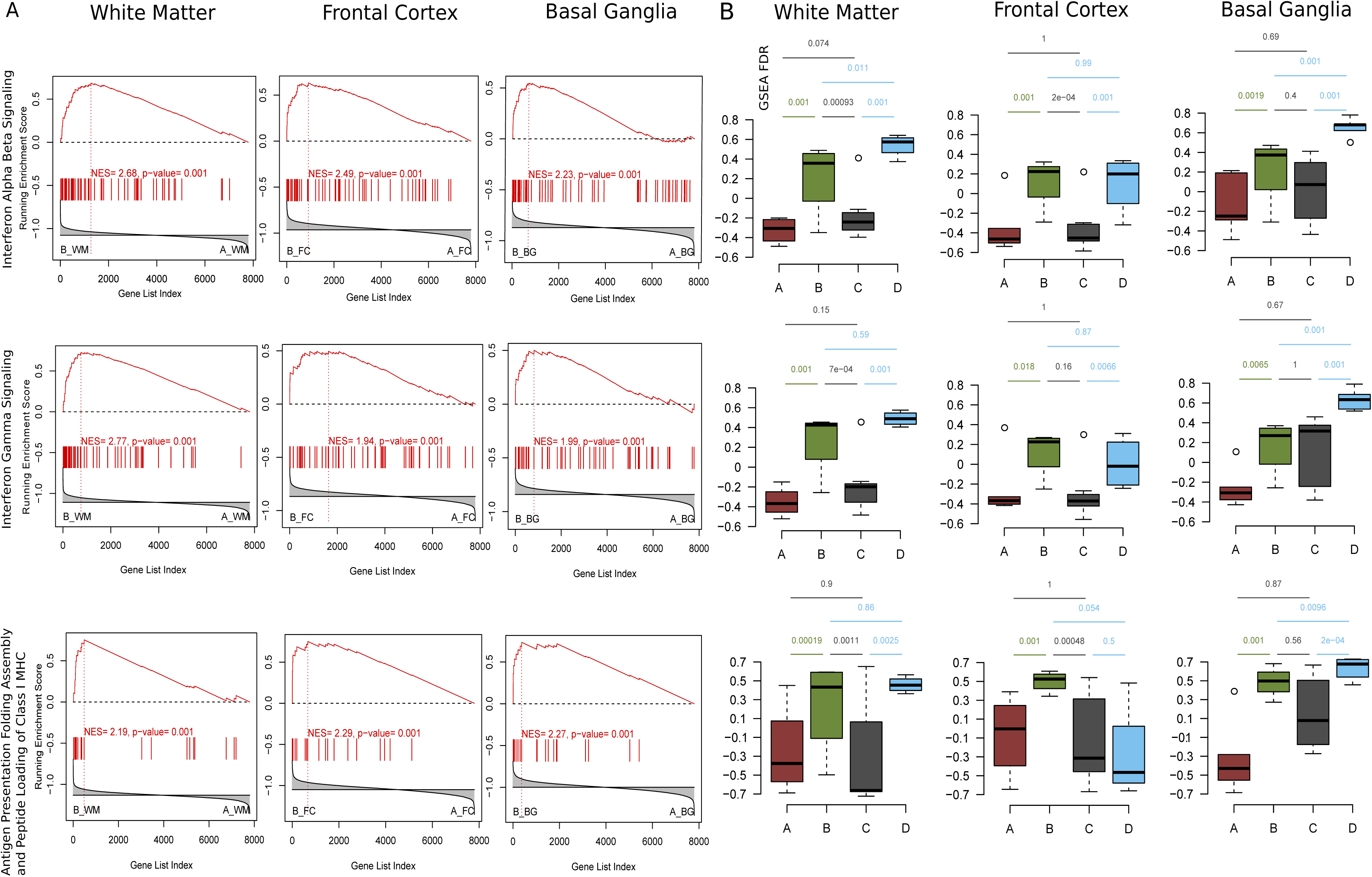
**Differential regulation of IFN-related pathways in the groups of the NNTC gene expression dataset. A) Gene expression evidence of interferon (IFN) activation HIV-infected patients without NCI.** The diagrams show GSEA plots for 3 pathways representative of IFN activation in HIV-infected patients without NCI (group B, left-hand side in the GSEA plot) as compared to uninfected controls (group A right-hand side). Each pathway was tested in each region independently. WM = White Matter; FC = Frontal Cortex; BG = Basal Ganglia. These pathways are indicative of type I IFN activation and include IFN-related genes in **Top**) INTERFERON ALPHA BETA SIGNALING, **Middle**) type II IFN activation (INTERFERON GAMMA SIGNALING), and **Bottom**) ANTIGEN PRESENTATION FOLDING ASSEMBLY AND PEPTIDE LOADING OF MHC CLASS I, a pathway involving several IFN-regulated MHC class I genes. Significant changes in the expression of the pathways is indicated by the asymmetric distribution of the genes in the geneset (vertical bars) and of the running enrichment score plot (ES) (19). Genes participating in the enrichment (on the left of the leading edge corresponding to the peak of the running enrichment in GSEA) are shown in Fig. 3. **B) Differential regulation of IFN-related pathways in the 4 groups of the NNTC gene expression dataset.** Significant activation of pathways related to both type I and type II IFN was seen in a brain region-specific pattern in HIV-infected patients without NCI (group B) and in patients with HIVE (group D). **Top**) INTERFERON ALPHA BETA SIGNALING in the white Matter, frontal cortex and basal ganglia; **Middle**) INTERFERON GAMMA SIGNALING; and **Bottom**) ANTIGEN PRESENTATION FOLDING ASSEMBLY AND PEPTIDE LOADING OF MHC CLASS I, in the same three regions. Each plot represents the pathway activity (computed as an enrichment score) in the 4 different phenotypes and is annotated with the FDR values of respective GSEA comparisons.

**Figure 3:**
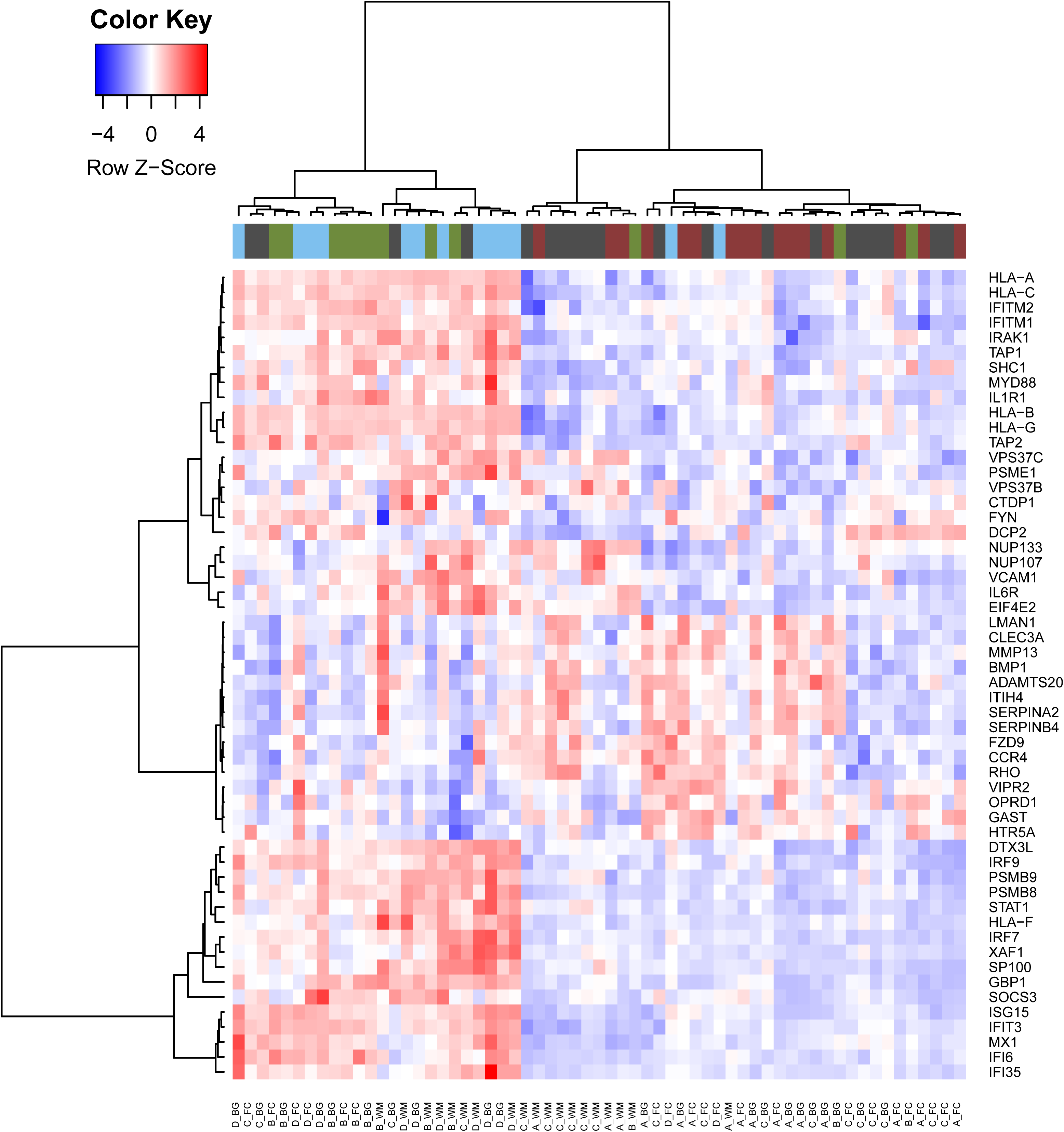
**Genes differentially expressed in HIV-infected patients without NCI.** The heatmap shows the genes in the leading edge of the pathways commonly dysregulated in all 3 brain regions in HIV-infected patients without NCI as compared to uninfected controls. We selected the 54 genes most differentially expressed (t-test, p-value < 0.01) among the list of 128 genes belonging to the leading edges of the significant pathways identified by the GSEA analysis in HIV-infected patients without NCI in comparison with uninfected controls.

We then looked at pathways specifically differentially regulated in one brain region as compared to the two other regions. To this end, we selected pathways enriched at FDR < 0.01 in one region and FDR > 0.25 in the other two. We observed 6 pathways meeting the criteria in the white matter, 4 in the prefrontal cortex and 2 in the basal ganglia. Pathways enriched in the white matter in the B-A comparison are indicative of immune activation, and complement induction (e.g., BIOCARTA COMP PATHWAY, BIOCARTA CTL PATHWAY, REACTOME IMMUNOREGULATORY INTERACTIONS BETWEEN A LYMPHOID AND A NON LYMPHOID CELL). We also observed increased expression of calpain-related genes (BIOCARTA UCALPAIN PATHWAY) in the prefrontal cortex and calpain-related and caspases-related genes in the basal ganglia (KEGG APOPTOSIS) as well as evidence of activation of the apoptosis-mediating p75 receptor (PID P75 NTR PATHWAY) and TNF-α signaling (PID TNF PATHWAY) in both prefrontal cortex and basal ganglia, indicative of tissue damage. Downregulation of genes related to neurotransmission was also evident in the prefrontal cortex and basal ganglia of patients with HIV but no NCI (group B) compared to control subjects (e.g., REACTOME LIGAND GATED ION CHANNEL TRANSPORT, KEGG NEUROACTIVE LIGAND RECEPTOR INTERACTION, Fig. 4), (Supplementary Table 2).

**Figure 4:**
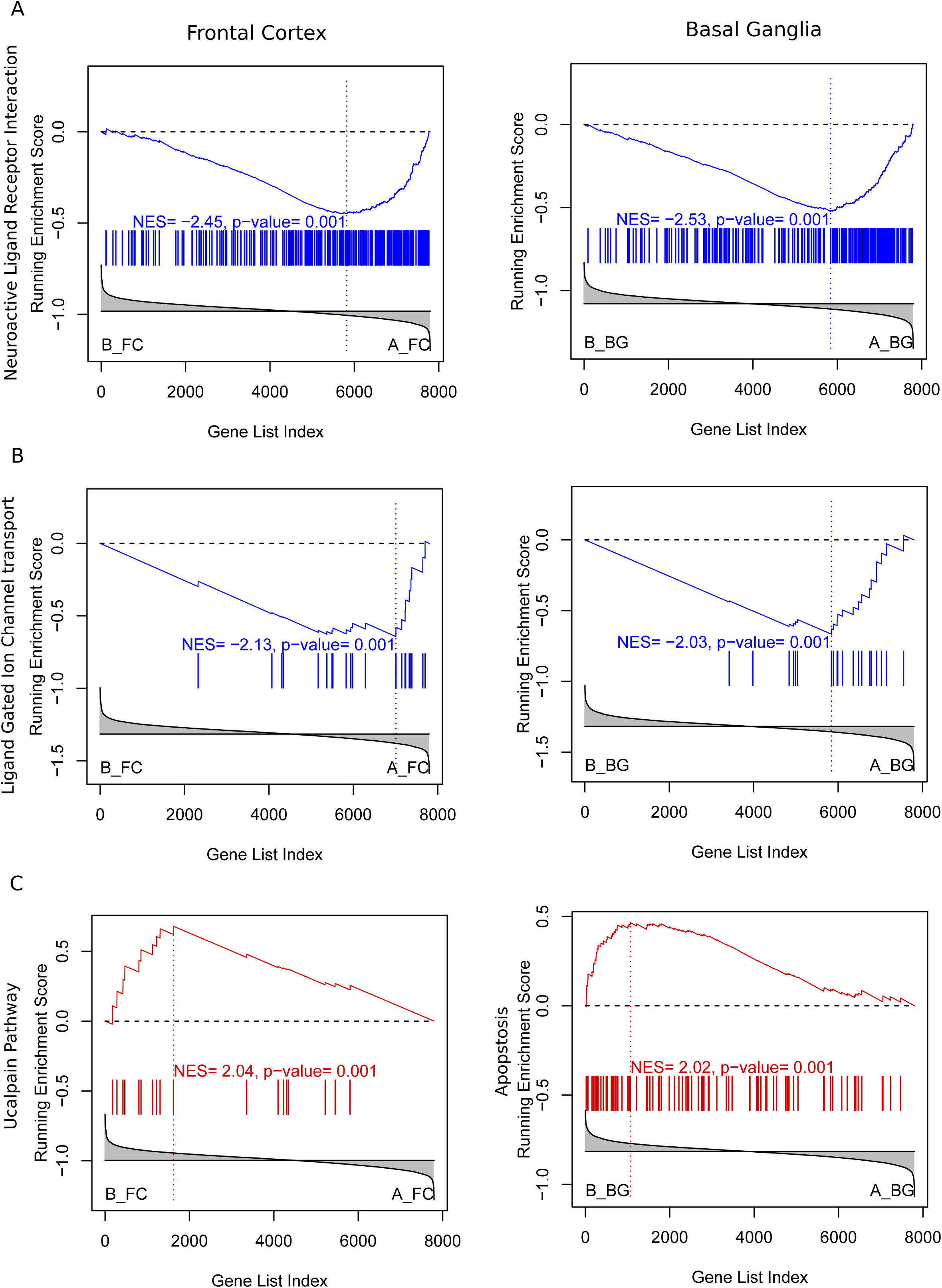
**Evidence of neuronal injury in HIV infected patients with HIV without NCI. A)** and **B)** Downregulation of genes related to neuronal transmission in patients with HIV without NCI (group B, left-hand side in the GSEA plot) vs. uninfected controls (Group A). **C**) Upregulation of apoptotic-related pathways in the frontal cortex and basal ganglia of HIV-infected patients without NCI (group B).

### Identification of pathways differentially regulated in HIV-infected patients with NCI without HIVE vs. uninfected controls (C-A comparison)

HIV-infected patients with NCI and no HIVE (group C), showed significant changes specific to the white matter compared to uninfected controls. Upregulated pathways are indicative of immune activation involving chemokine, cytokines and β-defensins induction (e.g., REACTOME CHEMOKINE RECEPTORS BIND CHEMOKINES; KEGG CYTOKINE CYTOKINE RECEPTOR INTERACTION, REACTOME BETA DEFENSINS, KEGG AUTOIMMUNE THYROID DISEASE, KEGG ALLOGRAFT REJECTION), oxidative stress and cytochrome P450 enzymes (KEGG DRUG METABOLISM CYTOCHROME P450, REACTOME BIOLOGICAL OXIDATIONS), matrix metalloproteases (MMPs) (NABA MATRISOME ASSOCIATED), and downregulation of genes related to RNA transcription and processing (e.g., REACTOME RNA POL II TRANSCRIPTION, KEGG SPLICEOSOME), (Supplementary Table 3), Fig. 5.

**Figure 5:**
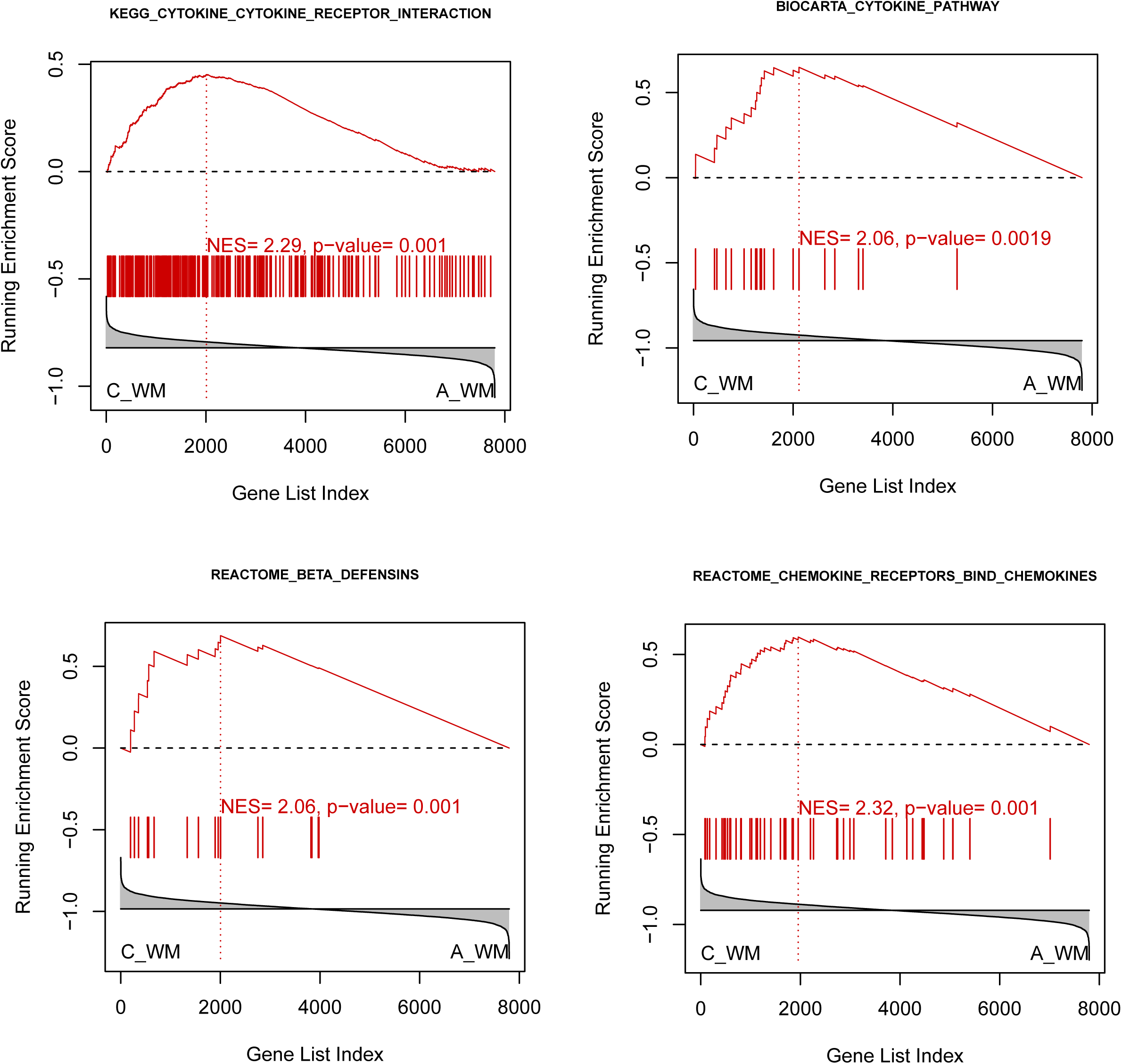
**White matter changes in HIV-infected patients with NCI without HIVE.** GSEA plots representative of induction of cytokines, chemokines and β-defensins in the HIV-infected patients with NCI without HIVE (group C in the NNTC gene expression dataset) as compared to uninfected controls.

### Identification of pathways differentially regulated between HIV-infected patients with NCI and no HIVE vs. HIV-infected without NCI (C-B comparison)

Two pathways were concordantly differentially regulated between groups B and C in all brain regions. These pathways are indicative of type I IFN activation in HIV-infected patients without NCI (group B), as indicated above. We also identified 47 pathways specifically differentially regulated in the prefrontal cortex. Among the pathways upregulated in group B as compared to C in the prefrontal cortex, were pathways indicative of tissue damage (e.g., REACTOME REGULATION OF APOPTOSIS), RNA transcription and processing (e.g., REACTOME METABOLISM OF RNA, KEGG RIBOSOME), and pathways related to protein degradation (e.g., KEGG PROTEASOME, REACTOME AUTODEGRADATION OF THE E3 UBIQUITIN LIGASE COP1, REACTOME APC C CDC20 MEDIATED DEGRADATION OF MITOTIC PROTEINS), (Supplementary Table 4).

### Identification of pathways differentially regulated between HIV-infected patients with HIVE vs. patients without NCI (D-B comparison)

Pathways dysregulated in all brain regions in HIVE (group D) as compared to patients with HIV without NCI (group B) are suggestive of disruption of protein folding, a mechanism of neurodegeneration (e.g., REACTOME POST CHAPERONIN TUBULIN FOLDING PATHWAY, REACTOME PREFOLDIN MEDIATED TRANSFER OF SUBSTRATE TO CCT TRIC). Other broadly dysregulated pathways in patients with HIVE (group D) as compared to patients with HIV without NCI (group B) indicate greater and broader activation of inflammatory and immune activation genes in group D as compared to group B. Genes regulated by both type I and type II IFN that were found activated in HIV infection without NCI (group B) as compared to uninfected controls, generally showed greater activation in patients with HIVE (group D). For instance, REACTOME INTERFERON GAMMA SIGNALING was increased in group D in basal ganglia in comparison to group B, and REACTOME INTERFERON ALPHA BETA SIGNALING was increased in group D in both white matter and basal ganglia in comparison to group B. All pathways specifically upregulated in basal ganglia in group D were involved in immune activation (e.g., REACTOME NUCLEAR EVENTS KINASE AND TRANSCRIPTION FACTOR ACTIVATION, REACTOME CYTOKINE SIGNALING IN IMMUNE SYSTEM, KEGG LEISHMANIA INFECTION, PID TCR PATHWAY).

Interestingly, the white matter in group D did not show any specific differentially regulated pathways in comparison to group B; conversely, the prefrontal cortex had 121 pathways and basal ganglia had 16 pathways significantly activated in group D in comparison to group B. Among the pathways upregulated in the prefrontal cortex in group D were pathways indicative of production of cytokine, chemokines and β-defensins (e.g., KEGG CYTOKINE CYTOKINE RECEPTOR INTERACTION, REACTOME CHEMOKINE RECEPTORS BIND CHEMOKINES, REACTOME BETA DEFENSINS). Pathways indicative of neurodegeneration were differentially regulated between D and B in frontal cortex including KEGG PARKINSONS DISEASE and KEGG HUNTINGTONS DISEASE (Fig. 6). These pathways include genes indicative of trophic interaction, protein misfolding and mitochondrial function. We also identified downregulated pathways related to mitochondria and energy metabolism were decreased in group D in all brain regions at FDR < 0.2 (e.g., REACTOME TCA CYCLE AND RESPIRATORY ELECTRON TRANSPORT, REACTOME PYRUVATE METABOLISM AND CITRIC ACID TCA CYCLE, REACTOME GLYCOLYSIS), (Supplementary Table 5).

**Figure 6:**
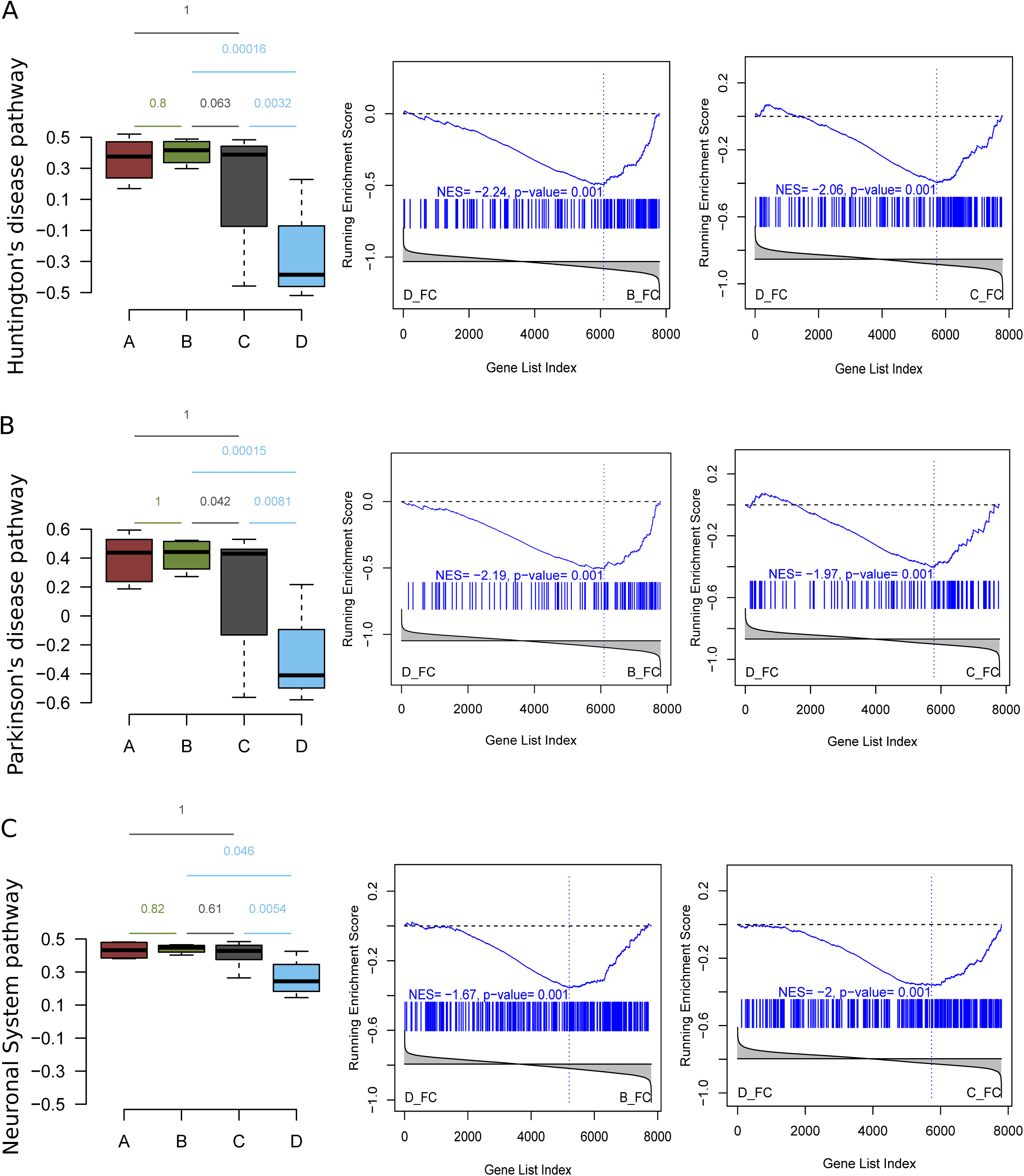
**Gene expression evidence of neurodegeneration in in Frontal Cortex of patients with HIVE. A)** Huntington’s disease-related pathways is downregulated in frontal cortex of patients with HIVE (group D) as compared to patients with NCI and no NIVE as well as with patients without NCI. Similarly, **(B)** and **(C)** show Parkinson’s disease and Neuronal System pathways respectively. Each row represents the pathway activity (computed as an enrichment score) in the 4 different phenotypes and is annotated with the FDR values of respective GSEA comparisons, followed by GSEA plots of the pathways in the group D vs group B and group D vs. group C comparisons in frontal cortex.

### Identification of pathways differentially regulated between HIV-infected patients with HIVE vs. patients with NCI and no HIVE (D-C comparison)

We identified 27 pathways concordantly differentially regulated at the C to D comparison. Seventeen pathways were upregulated in group D (HIVE) as compared to group C (NCI without HIVE) and largely reflected activation in HIVE of IFN response (e.g., REACTOME INTERFERON SIGNALING, REACTOME INTERFERON GAMMA SIGNALING, KEGG ANTIGEN PROCESSING AND PRESENTATION), immune activation and inflammatory cytokine signaling (e.g., REACTOME CYTOKINE SIGNALING IN IMMUNE SYSTEM, KEGG CYTOKINE CYTOKINE RECEPTOR INTERACTION, REACTOME INNATE IMMUNE SYSTEM), apoptosis (BIOCARTA DEATH PATHWAY), protein misfolding (REACTOME PREFOLDIN MEDIATED TRANSFER OF SUBSTRATE TO CCT TRIC), and HIV infection (PID HIV NEF PATHWAY, REACTOME LATE PHASE OF HIV LIFE CYCLE).

Ten pathways were downregulated in group D and included pathways related to translation and transcription, as seen in the A to B transition, likely reflecting transcriptional/translational dysregulations brought about by IFN activation (e.g., REACTOME TRANSPORT OF RIBONUCLEOPROTEINS INTO THE HOST NUCLEUS), HIV expression (REACTOME INTERACTIONS OF VPR WITH HOST CELLULAR PROTEINS, REACTOME LATE PHASE OF HIV LIFE CYCLE), impaired neuronal communication (REACTOME TRANSMISSION ACROSS CHEMICAL SYNAPSES), and energy metabolism (REACTOME CITRIC ACID CYCLE TCA CYCLE, REACTOME PYRUVATE METABOLISM AND CITRIC ACID TCA CYCLE).

Pathways related to neurodegenerative/neuronal pathways were differentially regulated in the prefrontal cortex and basal ganglia in patients with HIVE, including KEGG HUNTINGTONS DISEASE, KEGG PARKINSONS DISEASE, REACTOME NEURONAL SYSTEM (Fig. 6). A significant component of these pathways are genes involved in mitochondria function and energy metabolism. No pathways were specifically different in white matter between groups C and D, while 37 pathways were specific to the prefrontal cortex and 11 to basal ganglia. Several prefrontal cortex-specific pathways were downregulated in group D and included cell cycle regulation while basal ganglia pathways were related to translation/transcription and immune regulation (BIOCARTA D4GDI PATHWAY, BIOCARTA 41BB PATHWAY, PID CD8 TCR DOWNSTREAM PATHWAY, PID IL12 STAT4 PATHWAY), (Supplementary Table 6).

## Discussion

While dementia and HIV encephalitis are late consequences of HIV/AIDS, HIV enters the brain early after infection and remains in the brain throughout the course of infection. A considerable body of observations indicate that neuroinflammatory markers correlate with disease progression and the emergence of NCI in neuroAIDS (22–24). Proinflammatory cytokines and chemokines including IFN-α, TNF-α and CCL2 that are secreted by astrocytes and microglia have long been implicated in the pathogenesis of neuroAIDS (25–29). For instance, IFN-α in the cerebrospinal fluid has been observed to be higher in HAD compared with HIV-infected patients without HAD (25, 28, 30). Here, we show that HIV infection is associated with substantial dysregulations of gene expression related to immune activation before the onset of NCI and HIVE.

A primary finding in the present study is that we observed gene expression evidence of IFN induction in patients with HIV infection without NCI (group B) as well as in patients with HIVE (group D). Induction of both IFN type I and type II responsive genes was seen in patients with HIV infection without NCI (group B) in all brain regions studied. Among the genes differentially regulated within these pathways were IFN-responsive genes such as HLA-A, -B, -C, -G, -F, adhesion molecules such as VCAM-1, and ISG15 and IFI6 (31–33).

Chronic IFN expression is considered a key contributor to inflammation in neuroAIDS as well as a potential cause of NCI and depression vulnerability. However, data on the contribution of IFN activation to NCI are conflicting. Mice with transgene expression of IFN-α in astrocytes develop a dose-dependent inflammatory encephalopathy (34). Yet IFN-α transgenic expression in the central nervous system induced only mild effects in an egocentric spatial working memory test (35). However, the latter may also reflect compensatory changes as passively administered IFN-β impaired spatial memory in mice in another study (36). A recent study suggested a role for IFN-γ in shaping fronto-cortical connections and social behavior (37), which is consistent with a potential role of excessive IFN activation in the pathogenesis of NCI. In a recent large multi-center trial, depression was not significantly increased by IFNβ□treatment for multiple sclerosis (MS) (38). The early induction of IFN in patients of the NNTC dataset (patients with HIV without NCI, group B) is reminiscent of previous studies in which IFN induction was not closely correlated with NCI, e.g., (39), and raises the possibilities that either protracted IFN dysregulation may be required to produce NCI or that it may be a co-factor in NCI pathogenesis.

Also evident in HIV-infected patients without NCI was the activation of mechanisms indicative of tissue injury, such as expression of matrix metalloproteases (MMP) and complement-related genes in the white matter. MMP expression by HIV-1 infected monocytes and macrophages is recognized as a pathogenic mechanism in neuroAIDS (40). Elevated MMP levels can contribute to microglial activation, infiltrate through cleavage of adhesion molecules, neuronal and synaptic injury, as well as bloodbrain barrier disruption (41–44). MMP increases were present in the white matter in HIV-infected patients with NCI and no HIVE. In patients with HIVE, induction of MMPs was also evident in the prefrontal cortex and basal ganglia.

Another key finding in the study is that patients with NCI without HIVE (group C) in the NNTC cohort did not show significant activation of IFN, unlike patients in groups B and D. This discordant regulation of IFN signaling did not appear to be associated with antiretroviral therapy as patients with NCI and no HIVE include both patients treated with antiretrovirals and untreated patients. Conversely, patients with NCI without HIVE (group C) had increased expression of chemokines, cytokines and β-defensins in the white matter. Other pro-inflammatory markers were also concomitantly increased in the white matter of patients with NCI and no HIVE. Evidence of chemokine and cytokine expression were present in all HIV-infected groups in the study. β-defensins were induced also in patients with HIVE.

Chemokines have been implicated in impairing cognition, Alzheimer’s disease and depression as well as other psychiatric conditions (45). Increased immunoreactivity for MCP-2 was noted in MS lesions (46). A chemokine gene cluster has been associated with age of onset of Alzheimer’s (47). A higher level of CCL2 in CSF, and a CCL2 -2578G allele, have been associated with worse neurocognitive functioning in HIV (48). Animal studies, while scant, are consistent with a possible role for chemokines in NCI. For instance, chemokine signaling was increased by SIV infection and methamphetamine exposure in macaques (49, 50). Chemokines can induce changes leading to impaired hippocampal synaptic transmission, plasticity and memory (50, 51). Evidence also suggests a role for defensins in the chronic inflammation associated with degenerative brain diseases, and in particular Alzheimer’s disease (52, 53). Defensins-related pathways were also induced in HIVE, but showed no consistent regulation in HIV-infected patients with no NCI, suggesting a possible contribution to the pathogenesis of NCI.

In HIV without NCI, genes related to neurotransmission were also downregulated in the prefrontal cortex and basal ganglia while genes related to apoptosis, such as calpain-related mechanisms, which contribute to neurodegeneration in HIV (54), were induced in the basal ganglia and prefrontal cortex. Conversely, no pathways showed significant dysregulations in the prefrontal cortex and basal ganglia in HIV patients with NCI and no HIVE. In HIVE, multiple pathways indicative of impaired mitochondria and energy metabolism were differentially regulated. In NCI without HIVE, we observed increased expression of cytochrome P450 enzymes, which may indicate oxidative stress (55).

The anatomical distribution of the gene expression programs dysregulated in the NNTC dataset appears to reflect brain-region specific dynamics in neurological disease progression in HIV/AIDS. In particular, we observed some degree of white matter alteration of gene expression in all HIV-infected groups with and without NCI and HIVE. However, gene expression changes in patients with NCI without HIVE (group C) were localized to the white matter and had a specific gene expression profile. Lack of gene expression changes suggestive of neuronal injury in the prefrontal cortex and basal ganglia in patients with NCI without HIVE (group C) suggests that they may not be accompanied by significant neuronal atrophy, but that white matter pathology likely drives NCI in these patients. Prominent white matter gene expression changes were also present in HIVE, which was also characterized by considerable gene expression changes in the prefrontal cortex and basal ganglia. White matter damage correlating with the severity of cognitive manifestations has been observed since the early days of the HIV pandemic (8, 56). Evidence of white matter injury in HIV-infected patients with and without NCI is also demonstrated in recent imaging studies (57, 58). In addition to white matter changes, gene expression in HIVE was characterized by considerable changes in the prefrontal cortex and basal ganglia. This is also in apparent agreement with the association of NCI with progression of functional abnormalities involving the basal ganglia and the prefrontal cortex as well as with generalized white matter damage (56, 59–62).

The present study has several limitations. Primarily, the NNTC dataset groups are of small sample size that was further reduced as part of the quality control analysis. Larger studies will be needed to better understand the pathogenesis and progression of neurological disease and to adequately represent all possible variants of central nervous system disease. For instance, gene expression results of the group of HIV-infected patients with NCI and without HIVE raise several questions, including if this is a distinct nosologic variant of neuroAIDS or if it is a stage in the progression of HIV brain disease.

In conclusion, in the present study we explored patterns of gene expression dysregulation in patients in the NNTC neuroAIDS gene expression dataset. Results point to gene expression changes indicative of immune activation characterized by IFN and cytokine expression as well as evidence of neuronal suffering preceding NCI. Interestingly, the group of HIV-infected patients with NCI without HIVE showed a preeminently white matter dysfunction characterized by a distinct pattern of immune activation with low IFN. Larger studies are necessary to better understand the pathogenesis of neurological disease and its progression, to evaluate the impact of therapy on various HIV disease conditions, and to identify better therapeutic targets and strategies for NCI in HIV.

## Authors′ contributions

PPS and CL designed study, analyzed data PPS, VRC, EM, and CL interpreted results and wrote paper

## Acknowledgements

This is manuscript #29423 from The Scripps Research Institute. This work was supported by the National Institutes of Health grant numbers AA021667, DA041750, DA043268

